# Intrinsic dimensionality of single-cell transcriptomic data reveals potency landscapes during cell reprogramming

**DOI:** 10.1101/2025.07.21.665922

**Authors:** Maddalena Staiano, Niccolò Cirone, Marta Biondo, Matteo Osella, Antonio Scialdone

## Abstract

Cell potency, the ability of a cell to generate other cell types, is a fundamental property that drives development, regeneration, and reprogramming. Recent computational advances have enabled estimation of potency directly from single-cell RNA sequencing (scRNA-seq) data, with intrinsic dimensionality (ID) emerging as a promising, data-driven measure of transcriptional complexity linked to developmental potential. While ID offers an unbiased and interpretable framework, existing implementations have two main limitations: they have been applied in only a narrow range of biological contexts, and rely on clustering, which restricts resolution at the single-cell level. Here, we introduce IDEAS (Intrinsic Dimensionality Estimation Analysis of single-cell RNA sequencing data), a Python-based toolkit that computes both global and single-cell ID from scRNA-seq data, enabling potency scoring without requiring predefined clusters. IDEAS extends ID-based potency estimation to single cells, showing its usefulness in a new biological setting. It offers a clear and reliable way to study cell plasticity, simplifies ID score calculation, and makes the method easier to use across different biological systems, speeding up research on this approach to measuring cell potency. We apply IDEAS to multiple datasets of cellular reprogramming, a dynamic and heterogeneous process in which cells transition between identities. Our analysis reveals that ID scores effectively capture changes in potency, identify partially reprogrammed intermediates, and delineate alternative reprogramming trajectories.

## 1 Introduction

Cells exhibit a remarkable diversity of types, each defined by distinct functions and varying capacities to alter their identity and differentiate. Among these, stem cells stand out for their ability to give rise to all other cell types in the body. This capacity is referred to as *cell potency*, which exists along a spectrum. Terminally differentiated cells, with highly stable identities, are considered to have the lowest potency, while pluripotent stem cells, capable of differentiating into any cell type, represent the highest potency level. This developmental potential is a fundamental property that underpins essential biological processes in multicellular organisms, such as embryogenesis and blood cell differentiation [1, 2, 3, 4]. Its medical significance is particularly profound, as it forms the basis for advances in regenerative medicine, where researchers seek to manipulate cells to modify their potency and identity [5, 6, 7, 8, 9].

Cell potency is typically evaluated using a combination of experimental approaches. Functional *in vitro* differentiation assays assess a cell’s capacity to generate derivatives of one or more germ layers, often employing directed differentiation protocols followed by phenotypic and molecular characterization of the resulting lineages [10, 11, 12]. *In vivo* assays, such as teratoma formation assays for pluripotent cells [13] or chimera contribution analyses in animal models [14], determine the ability of a stem cell to integrate into host tissues and contribute to normal development. Additionally, *in vivo* engraftment assays are used to test the regenerative potential and lineage output of tissue-specific stem cells [15, 16]. In addition to these functional assessments, marker-based assays, including quantitative RT-PCR, flow cytometry, and immunochemistry, are employed to quantify the expression of lineage- or potency-associated markers, providing molecular signatures indicative of the identity and developmental potential of a cell [17, 18]. While experimental approaches can be accurate, they are often expensive and labor-intensive. Moreover, marker-based assessments rely on prior knowledge and may lack flexibility.

To address these limitations, several computational methods have been proposed to estimate cell potency from single-cell RNA-seq (scRNA-seq) data relying on measures of transcriptional diversity and entropy. For example, these include SCENT, which incorporates protein-protein interaction networks [19]; SLICE, which leverages Gene Ontology annotations [20]; and CytoTRACE, which infers potency based on the number of expressed genes per cell [21]. More recently, a geometry-based approach has been introduced that estimates cell potency via the intrinsic dimensionality (ID) of scRNA-seq data [22], offering an alternative that does not depend on prior biological knowledge, such as marker genes, or external annotations and databases. Although this method has shown promise in developmental biology, its current implementation relies on cell clusters, providing a score for cell populations and thus limiting its resolution and single-cell applicability [22].

Building on this foundation, here we present IDEAS (Intrinsic Dimensionality Estimation Analysis of single-cell RNA sequencing data), a Python-based toolkit that integrates with established frameworks for scRNA-seq data analysis like Scanpy [23]. IDEAS extends previous methodologies by enabling both global and local ID computation from scRNA-seq data, the latter assigning an ID-based pluripotency score to each individual cell, thus eliminating the need for predefined cell populations. We demonstrate the utility of IDEAS by applying it to several datasets in the context of cellular reprogramming, a domain where the use of intrinsic dimensionality has not been previously explored. Cellular reprogramming is a process during which cells are induced to change their identity, e.g., to revert to a stem cell-like state [24] [25] or to switch to another fully differentiated cell type (trans-differentiation or direct reprogramming) [26, 27, 28]. Reprogramming has been extensively studied in the last twenty years as an extremely relevant advancement in regenerative medicine and stem cell therapies [29, 30]. Despite the increasing efforts in this direction, many aspects of cellular reprogramming remain to be understood, such as its low efficiency and asynchronous nature [31, 32]. Another critical aspect is that cells from the initial population do not undergo reprogramming in a deterministic manner: instead, a multitude of possible developmental pathways often arises from the same cell type [33], including pathways that terminate in only partially reprogrammed cells [34, 35, 36]. The identification and classification of such pathways can be very challenging.

Using IDEAS, we show that intrinsic dimensionality correlates with cell potency during reprogramming, allowing us to reconstruct and classify reprogramming trajectories, including direct and indirect transitions, and to detect intermediate, partially reprogrammed states. Through both cluster-based and single-cell analyses, IDEAS offers a robust and user-friendly framework for unbiased potency estimation via intrinsic dimension computation, opening new avenues for dissecting developmental processes and cellular plasticity.

## 2 Results

### 2.1 Overview of the method

IDEAS is a user-friendly Python toolkit designed for estimating the intrinsic dimension (ID) of single-cell RNA sequencing (scRNA-seq) datasets. It integrates multiple ID estimation methods along with all essential preliminary steps of the analysis, as illustrated in Figure 1. Below, we provide an overview of each step in the workflow and summarize the ID estimation algorithms currently implemented in IDEAS.

**Figure 1:**
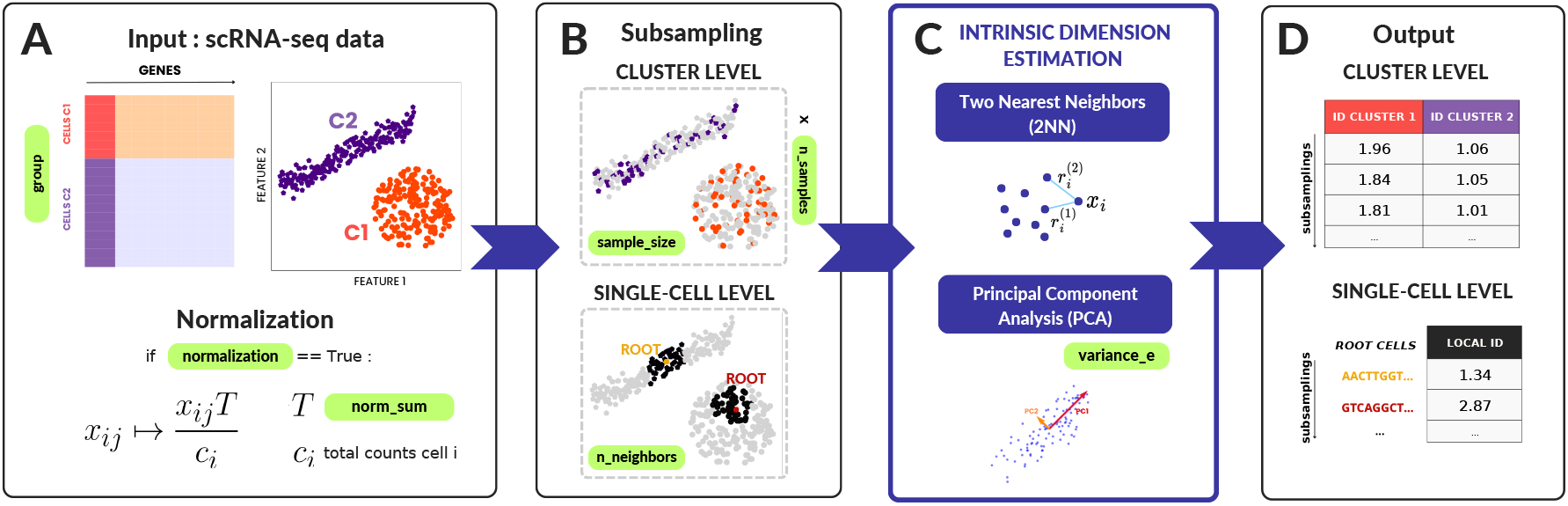
Illustration of the IDEAS workflow: **A)** IDEAS takes in input the scRNA-seq count matrix as an Annotated Data Object with cluster annotations (or day, cell type, etc.) for each cell. Single cells can be represented as data points in the gene expression space. Cell counts are normalized to a chosen target sum. **B)** Subsampling is applied to select an equal number of cells for each estimation. Two subsampling strategies are available: the population-level (cluster-level) approach selects an equal number of cells per group via random sampling; the single cell (local) approach selects nearest neighbors around a set of root cells **C)** Intrinsic dimensionality (ID) is estimated for each subsampling using one of two methods: Two Nearest Neighbors (TWO-NN) or Principal Component Analysis (PCA). **D)** The output is a Pandas DataFrame containing the estimated ID values. Each row corresponds to a subsampling. In the cluster-level approach, rows represent random subsamplings, and columns correspond to cell groups. In the single-cell approach, rows correspond to local subsamplings and are indexed by root cell barcodes, allowing direct mapping between cells and their local ID estimates.

#### Input

The input required by IDEAS is an Annotated Data Object, where genes correspond to variables (var) and cells correspond to observations (obs). With the `group` parameter, the user can specify the groups of cells for which the ID is computed and then compared (Figure 1A); if group is not specified, the ID is computed by default on the whole dataset.

#### Normalization

As it is standard practice in most scRNA-seq data analyses [37], ID estimation requires a normalization step and, possibly, the removal of batch effects [38]. IDEAS can normalize RNA counts for sequencing depth by rescaling them to a specified target value (set using the parameter `norm_sum`), provided that the parameter `normalization` is set to *True* (Figure 1A).

#### Intrinsic Dimension Estimation

The core of IDEAS consists in the evaluation of the intrinsic dimension [39]. In the context of data science, there is a wide variety of intrinsic dimension estimators [40, 41, 42, 43, 44], each based on different assumptions about the data. In IDEAS, the parameter method allows the user to choose between a fractal estimator (TWO-NN) [43, 44] and a PCA-based one [45, 41] (Fig. 1C). Additional estimators can be easily added in future versions of the package by passing custom NumPy [46] functions to the method parameter. In the present work, we report results obtained with TWO-NN, which has been selected as the default method in IDEAS due to its higher robustness and capacity to capture nonlinearities (further details in Supplementary Material).

#### Population-based versus single-cell potency estimation

Setting the `method` parameter to either *2nn* or *pca* enables the estimation of ID of predefined cell populations, as specified by the parameter group mentioned above. In general, these ID estimators bare functions of the sample size, i.e. the number of points (cells) used [22]. Therefore, since groups typically differ in the number of cells, their ID values cannot be directly compared. To compensate for this effect, IDEAS randomly sub-samples a fixed number of cells (`sample_size` parameter, with the default value corresponding to 80% of the smallest group size) from each group. The random subsampling is performed multiple times (`n_samples` parameter, 30 by default), and the average and standard deviation are calculated. In this approach, a single mean ID value is assigned to each pre-defined group of cells (e.g., via clustering).

However, this strategy may overlook intra-cluster heterogeneity, which can be crucial when investigating cell-to-cell variability, and the user could be interested in an estimate of cell potency for individual cells. To address this limitation, IDEAS offers an alternative method based on local sub-sampling. This approach assigns an ID value to each individual cell by considering its local neighborhood, thereby achieving single-cell resolution and enabling a more nuanced exploration of cellular heterogeneity [47]. The single-cell mode can be activated by setting the `method` parameter to *local 2nn* or *local pca*. In this case, a set of ‘root’ cells is selected through the parameter roots, by default set to the entire dataset. For each root cell, the ID is estimated on its neighborhood, i.e. the set of its *N* nearest neighbors defined with a Euclidean metric. The parameter `n_neighbors` allows to set the resolution *N*, by default set to 10% of the total cell count. A too local choice of *N* can make the ID estimation affected by sampling noise, but is better suited to keep into account data heterogeneity and non-linear correlations. Conversely, larger resolutions lead to more robust outcomes, but are sensitive to large-scale patterns such as the simultaneous presence of multiple cell types [22]. A grid search over multiple values of *N* is recommended to identify the optimal resolution for a given dataset (more details are given in Supplementary Material).

Ultimately, the population-based ID measurement is computationally more efficient and suitable for pipelines aimed at capturing global differences in the cell potency between well-defined populations, such as cells from different time points, cell types or biological replicates. When, instead, no clear grouping structure emerges from the data, or when class sizes are very imbalanced, the single-cell measure of ID can give meaningful insights on local and continuous variations of potency in single cells. Further insights on the advantages of this local approach are discussed in Section 2.3 and Supplementary Material.

#### Output

The output of IDEAS is a pandas DataFrame containing the estimated values of ID (Figure 1D), where each row (index of the DataFrame) represents a subsampling. When using the population-based approach (i.e., the *2nn* or *pca* methods), rows correspond to random subsamplings and columns to groups of cells. Setting the parameter `full_output` to *False*, IDEAS retrieves only the mean and standard deviation of the ID estimates, providing a simplified output that can be plotted as a single data point (with error bar) for each cell population. When instead the local subsampling is used (i.e., with the *local 2nn* or *local pca* methods), each row index in the DataFrame corresponds to the barcode of the ‘root’ cell used for the local ID estimation. This enables a direct mapping between single cells and their local ID.

The absolute values of ID are not inherently meaningful, as they depend on the specific subsampling size used [22]. Instead, ID should be interpreted as a relative measure, suitable for capturing trends in cell potency and for comparing groups or single cells. Therefore, to facilitate interpretation, we rescaled the ID values to the [0,1] range in subsequent analyses and refer to this normalized measure as the *ID-score* (see Section 4). This rescaling can be enabled by setting the `id_score` parameter to *True* (See Section 4.2).

### 2.2 Using IDEAS on scRNA-seq data from cell reprogramming experiments

In this study, we use IDEAS to investigate how the ID of cell transcriptomes varies during cell reprogramming. Many critical aspects of this process are still unclear, including the identity of emerging cell states and the developmental trajectories that lead to successful or failed reprogramming. Cellular reprogramming is generally conceptualized as an induced shift in cell identity that can be represented as a trajectory on the epigenetic landscape. For example, during the reprogramming of somatic cells into iPSCs, the initial cell population undergoes a de-differentiation process, resulting in a final population with greater cell potency than the original. According to the epigenetic landscape metaphor, differentiated cells, which are initially located in a local minimum of the rugged landscape, are expected to “roll up” toward higher-energy states, corresponding to increased potency [1]. We first focus on this reprogramming process from somatic cells to iPSCs (see Section 2.2.1), as it provides a clear expectation for the cell potency trend that can be used to evaluate the reliability of the ID estimations provided by IDEAS.

Afterwards, we analyze datasets generated from direct reprogramming or transdifferentiation experiments (Section 2.2.2). In this context, somatic cells are directly converted into a different somatic cell type without transitioning through an intermediate pluripotent state. Using again the landscape analogy, this corresponds to a trajectory from one local minimum to another, without necessarily passing through high potency states. How this trajectory evolves on the landscape is difficult to determine. Consequently, unlike the first type of reprogramming, there is no clear expectation about the trend in cell potency during transdifferentiation.

Finally, we analyze a dataset capturing human fibroblast reprogramming to iPSCs, in which multiple divergent trajectories emerge, including both successful and failed routes towards pluripotency, highlithing the potential of ID to discriminate them (Section 2.2.3).

By analyzing these scenarios, we aim to show how IDEAS can reveal the trends of transcriptional trajectories and provide meaningful insight into cellular identity transitions.

#### 2.2.1 IDEAS reveals intrinsic dimension trends in single-cell data during mouse iPSCs generation

The reprogramming of somatic cells into induced pluripotent stem cells (iPSCs) offers a powerful model for studying cellular plasticity and the mechanisms of transcriptional and epigenetic remodeling. Mouse embryonic fibroblasts (MEFs) offer several practical advantages for studying cellular reprogramming to iPSCs, including a high proliferation rate, genetic stability, and responsiveness to reprogramming factors. Notably, many MEF-based reprogramming datasetes include longitudinal sampling, which makes them particularly suitable for analyzing reprogramming dynamics and evaluating computational methods. A common approach to tracking the differentiation state of MEFs during their transition into iPSCs involves monitoring a small set of pre-selected marker genes [48].

In [49], the transcriptional changes occurring during MEFs reprogramming into iPCSs via overexpression of the OSKM reprogramming factors (Oct4, Sox2, Klf4, and c-Myc) are tracked using scRNA-seq. The experiments were carried out under two different culture conditions: with fetal bovine serum (FBS) and with a combination of epigenomic modifiers and signaling molecules referred to A2S (ascorbic acid, 2i, SGC), designed to increase the efficiency of reprogramming.

Single cells were sampled and processed at different time points during the reprogramming protocol, and we analyzed the dynamics of ID along the time course separately for each experimental condition. Given the stochastic and asynchronous nature of the reprogramming process, each time point exhibits varying degrees of transcriptional heterogeneity. This could be due to the variability in the rates of de-differentiation of individual cells [50, 34] and the possibility of branching events into alternative cell types [51, 52, 53, 54]. To mitigate these potential confounding factors, we first identified transcriptional clusters using the Leiden algorithm [55]. In both culture conditions, the starting MEF population and the pluripotent mESC population predominantly form distinct clusters (Figure 2C,F), while a subset of cells from day 12 clusters with the mESCs, suggesting that these cells may have undergone successful reprogramming. Instead, cells from intermediate time points appeared more dispersed across multiple clusters. In the FBS condition, cells from intermediate days grouped into three clusters with varying proportions. On the contrary, in the A2S condition, the time mixing within the clusters was less pronounced, with cells from day 2 clustered into two distinct populations (Figure 2C,F).

**Figure 2:**
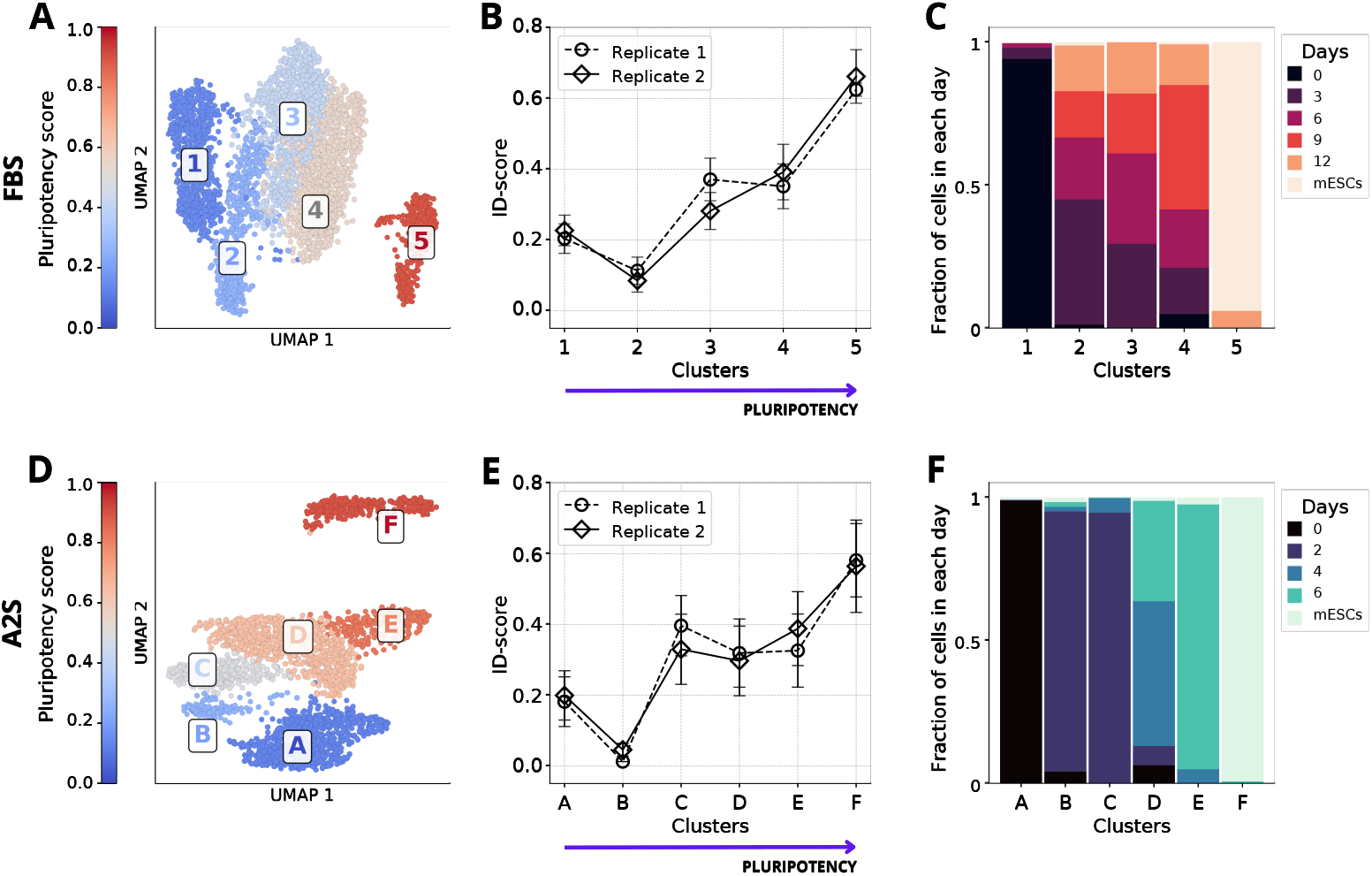
The intrinsic dimension increases with the potency level during reprogramming. **A**,**D)** UMAP projections of data from [49] in the FBS (panel A) and the A2S (panel D) conditions. Clusters are colored according to their pluripotency score (see Methods). **B**,**E)** ID-scores across clusters in the FBS (panel B) and A2S (panel E) conditions in two replicates (continous and dashed black lines). Each value represents the mean of the ID estimations computed over 200 random sub-samples and the error bars indicate the standard deviation. The clusters are sorted on the x-axis according to their pluripotency score. During sub-sampling, we used a sample size of 140 and 60 for data from the FBS and the A2S conditions respectively. **C**,**F)** Barplots illustrating the day-wise distribution of cells across clusters under the FBS (Panel C) and A2S (Panel F) conditions. Each plot is normalized to the total number of cells per day, allowing comparison of relative cluster proportions over time.

After identifying cell populations through clustering, we defined a pluripotency score based on the mean expression level of differentiation-related genes identified by the authors [49] (See Section 4.3.1). Figure 2A,D show a UMAP projection [56] of the data, where cells are colored by cluster and the color gradient reflects the average pluripotency score of each cluster. As expected, clusters that include cells from earlier time points have a lower pluripotency score compared to clusters that are enriched in cells from later time points. This biologically-informed score enabled us to order the clusters according to their expected potency levels. We further validated this ordering using a pseudotime analysis [57] (see Section 4.3.3 of Methods).

We then applied IDEAS to perform a fully unbiased estimation of cell potency using the ID. The resulting ID-scores show a positive correlation with the pluripotency scores of the clusters (Figure 2B,E). This trend is consistent across replicates and culture conditions. These results suggest that IDEAS can effectively capture the increase in cell potency during the reprogramming of MEFs into iPSCs without relying on predefined gene sets, thus providing a general and unbiased metric of the developmental potential.

#### 2.2.2 IDEAS reveals divergent intrinsic dimension trends in iPSC reprogramming versus transdifferentiation

As discussed above, reprogramming can give rise to several distinct cell fates with different levels of potency. These alternative fates often manifest as transcriptional trajectories in scRNA-seq data, which can be identified using specialized computational tools [58]. The states that cells transition through can be characterized by analyzing the trends of marker genes along the trajectories. However, this process is not always straightforward, as marker genes may be suboptimal or, in some cases, entirely unknown. In this section and the next, we leverage IDEAS to predict the potency level of cell states along different trajectories in a manner that is completely agnostic to marker genes. This analysis was conducted on two scRNA-seq datasets, each capturing reprogramming pathways of varying complexity.

In the first scRNA-seq dataset we analyzed, Francesconi et al [59] employed C/EBP*α* and OSKM induction to convert the initial population of murine pre-B cells into either induced macrophages (iMac) or iPSCs. In particular, pre-B cells were incubated with E2 for 18 hours to transiently activate C/EBP*α*, generating a ‘poised state’. After this timepoint, cells were divided in distinguished samples and treated with either C/EBP*α* or OSKM induction to generate different cell fates. In this context, we refer to the first process as transdifferentiation, which leads to the formation of an iMac population, and the second as reprogramming, which results in an iPSC sample (Figure 3A).

**Figure 3:**
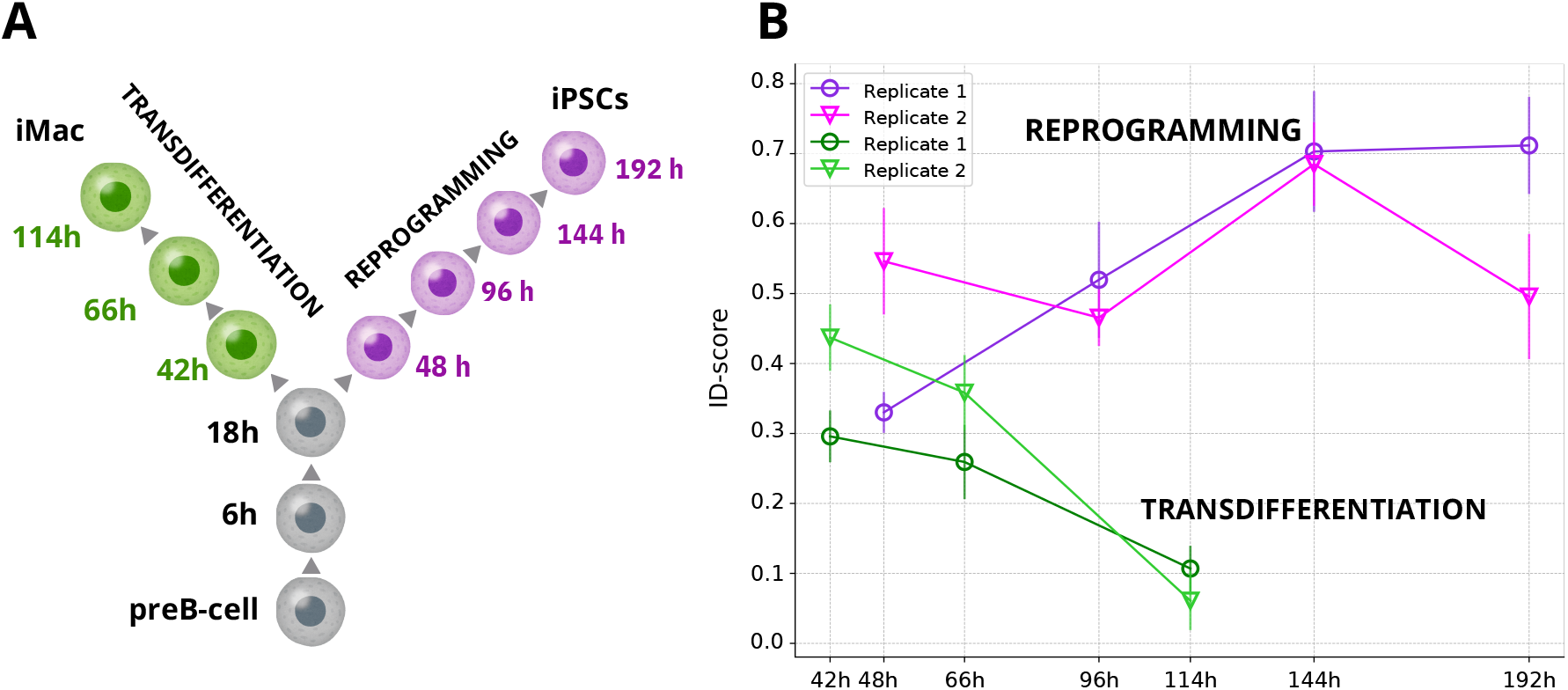
Intrinsic dimension reveals divergent trends during reprogramming and transdifferentiation -. **A)** Schematic representation of the experimental design in [59] with different sampling times in the two branches. **B)** Time-resolved ID trends reconstruct the bifurcation of the two trajectories. The ID score was estimated with a sample size of 180 cells and 1000 sub-samples per cluster.

Figure 3B presents the temporal trends of ID-scores obtained using IDEAS. These trends vary between the two trajectories leading to either iMac or iPSC states. Specifically, the ID-score decreases along the transdifferentiation trajectory as cells progress towards iMac differentiation. In contrast, the ID score for cells reprogrammed into iPSC either increases (in replicate 1) or oscillates (in replicate 2), while consistently remaining higher than the ID score observed along the iMac trajectory. This pattern aligns with the expected differences in cell potency between the two pathways.

It is important to note that these trajectories begin from an 18-hour “poised state” representing an intermediate stage of differentiation. As a result, the observed trends reflect changes in cell potency from this intermediate state onward. To gain deeper insights into how potency evolves across these trajectories, further studies are required, including more frequent sampling from initial to final populations and analyses of larger cell numbers.

Overall, these findings show that the ID-score estimated using IDEAS effectively captures the bifurcation event that distinguishes the transdifferentiation and reprogramming trajectories.

#### 2.2.3 The ID-score distinguishes multiple transcriptional trajectories towards divergent cell fates during reprogramming

During reprogramming, multiple cell fates can emerge. Using the data in [34], we tested whether the ID-score can capture cell potency trends in a more complex scenario, where a large number of cell states and multiple trajectories are present. In [34], the authors used scRNA-seq to investigate the dynamics of cellular reprogramming from human fibroblasts to induced pluripotent stem cells (iPSCs) by inferring transcriptional trajectories with the PAGA algorithm [60]. By analyzing the expression patterns of a subset of well-established marker genes, they identified the trajectories illustrated in Figure 4A as well as the associated cell types:

**Figure 4:**
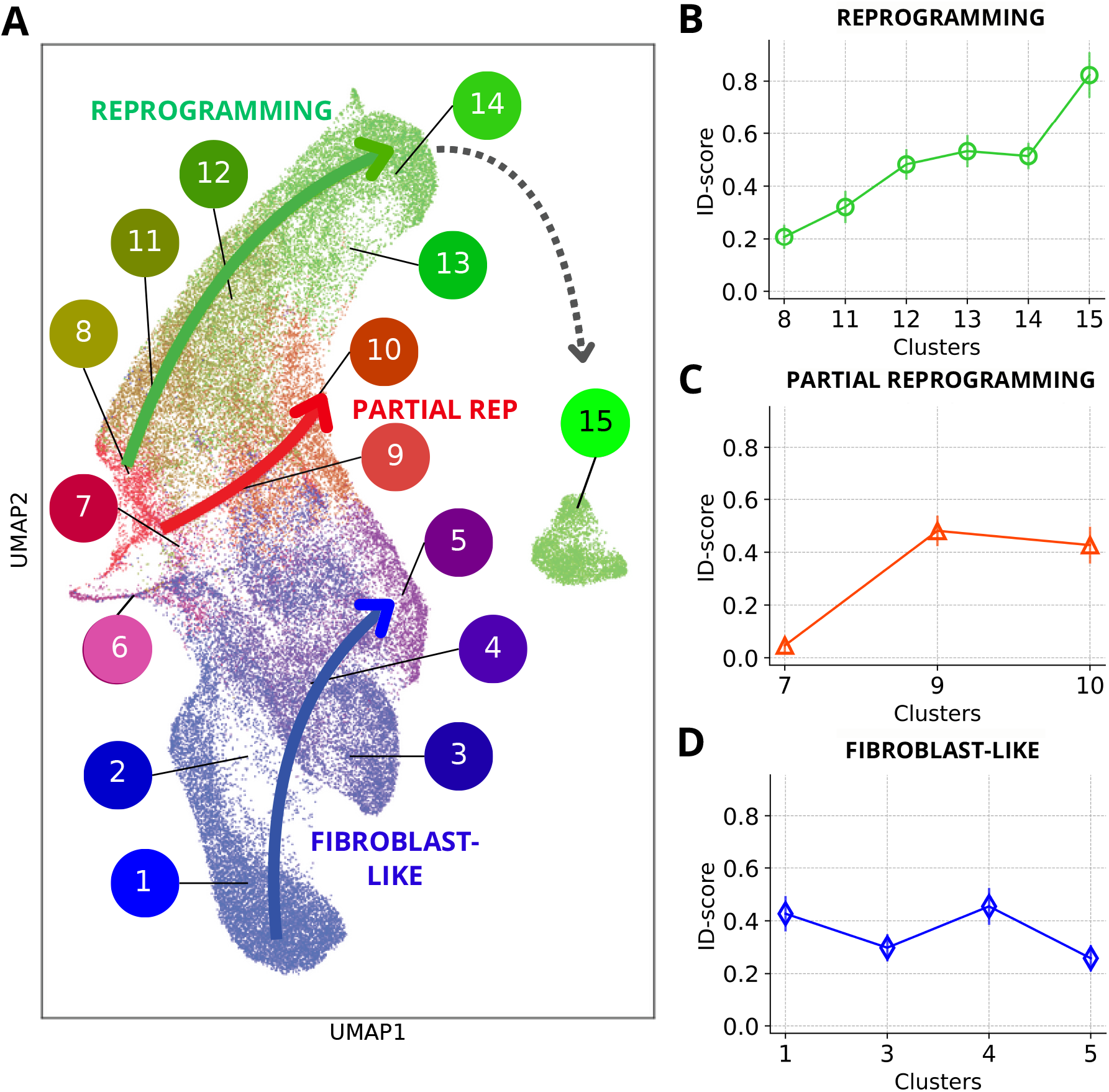
Intrinsic dimension can distinguish trajectories emerging during cell reprogramming -. **A)** UMAP projection [56] of cells grouped in clusters and trajectories obtained in [34] using the PAGA algorithm [60]. **B-D)** ID scores computed for clusters along the reprogramming (panel B), the partial reprogramming (panel C) and the fibroblast-like trajectory (panel D). The ID was estimated with a sample size of 418 cells and 50 sub-samples per cluster. Cluster 2 was removed from the analysis due to the low number of cells.

- *Fibroblast-like trajectory* : This trajectory encompasses clusters 1-2-3-4-5 and includes cells that retain fibroblast identity throughout the reprogramming process.
- *Partial reprogramming trajectory* : Clusters 7-9-10 belong to this trajectory, comprising cells that initiate reprogramming but fail to activate endogenous OCT4 expression, distinguishing them from pre-iPSC cells.
- *Reprogramming trajectory* : This trajectory, involving clusters 8-11-12-13-14-15, includes cells that successfully complete reprogramming to iPSCs.
- *Keratinocyte-like population*: Cluster 6 represents cells that do not belong to any of the trajectories mentioned above and exhibit keratinocyte-like characteristics.

Figure 4A displays a UMAP projection of the dataset, with cells colored by clusters and arrows illustrating the identified trajectories.

We applied IDEAS to this dataset to test whether the ID-score could track changes in potency along distinct transcriptional trajectories. The ID-score trends corresponding to the three primary trajectories are plotted in Figure 4B,D. Notably, the ID-score shows a pronounced increasing trend exclusively in the *reprogramming trajectory*, consistent with the rise in potency as cells reprogram to iPSCs already observed in the previous datasets. In contrast, the *partial reprogramming trajectory* exhibits an increase only between the first two clusters, reflecting a failure to fully activate key pluripotency markers. Finally, the *fibroblast-like* clusters only display fluctuations in the ID-score without a discernible trend. Cells at the endpoint have an ID score even slightly lower than the initial one, a result that again aligns with their persistent fibroblast marker expression [34]. Moreover, the ID-score of the *keratinocyte-like* population was the lowest among all clusters, consistent with its identity as a more differentiated cell state.

These findings demonstrate that the ID-score can distinguish trajectories based on cellular potency dynamics, supporting IDEAS as a powerful and general-purpose tool to investigate cell fate transitions.

### 2.3 Intrinsic dimension estimation on local neighborhood of cells as a single-cell potency measure

In the previous sections, we computed the ID of groups of cells primarily identified through clustering. However, this approach inherently depends on the number and composition of clusters, which remain somewhat arbitrary due to the ill-posed nature of clustering. While robustness analyses and manual annotation using marker genes can help validate the choice of clusters, the final partitioning of the data remains influenced by the selected clustering procedure criteria and priors. Moreover, in some biological processes, cell identity, and thus also potency, may be more accurately represented as a continuous spectrum rather than discrete categories defined by hard cluster membership. To circumvent these limitations, it is useful to perform analyses that do not rely on predefined clusters.

A possibility is to estimate ID directly at the level of local cell neighborhoods, for instance, by leveraging k-nearest neighbors (kNN) graphs. Although some parameters have to be selected, such as the number of neighbors, this strategy does not require clustering and provides a local ID estimation centered on each single cell. Therefore, we implemented a local version of the ID estimation, where the ID is computed using the kNN neighborhood for each cell in the dataset. This localized ID estimation serves as a more granular measure of cellular potency, and provides a score at the single-cell level.

To validate this approach, we first applied the local ID estimation to datasets analyzed in previous sections. Reproducing the analysis from Section 2.2.1 at single-cell resolution, we observed a strong positive correlation between the local ID of single cells and their corresponding pluripotency score (defined as Eq.2), for both FBS and A2S culture conditions (Figure 5 A,B). Notably, this correlation persists even when varying the number of nearest neighbors, indicating that our approach is robust to parameter selection (Supplementary Figure 4). Coloring cells by cluster revealed that local ID values are broadly consistent with cluster-level estimates (Figure 5). Finally, we confirmed a similarly strong correlation between local ID and pseudotime at the single-cell level, further supporting the validity of this local potency measure (Supplementary Figure 3).

**Figure 5:**
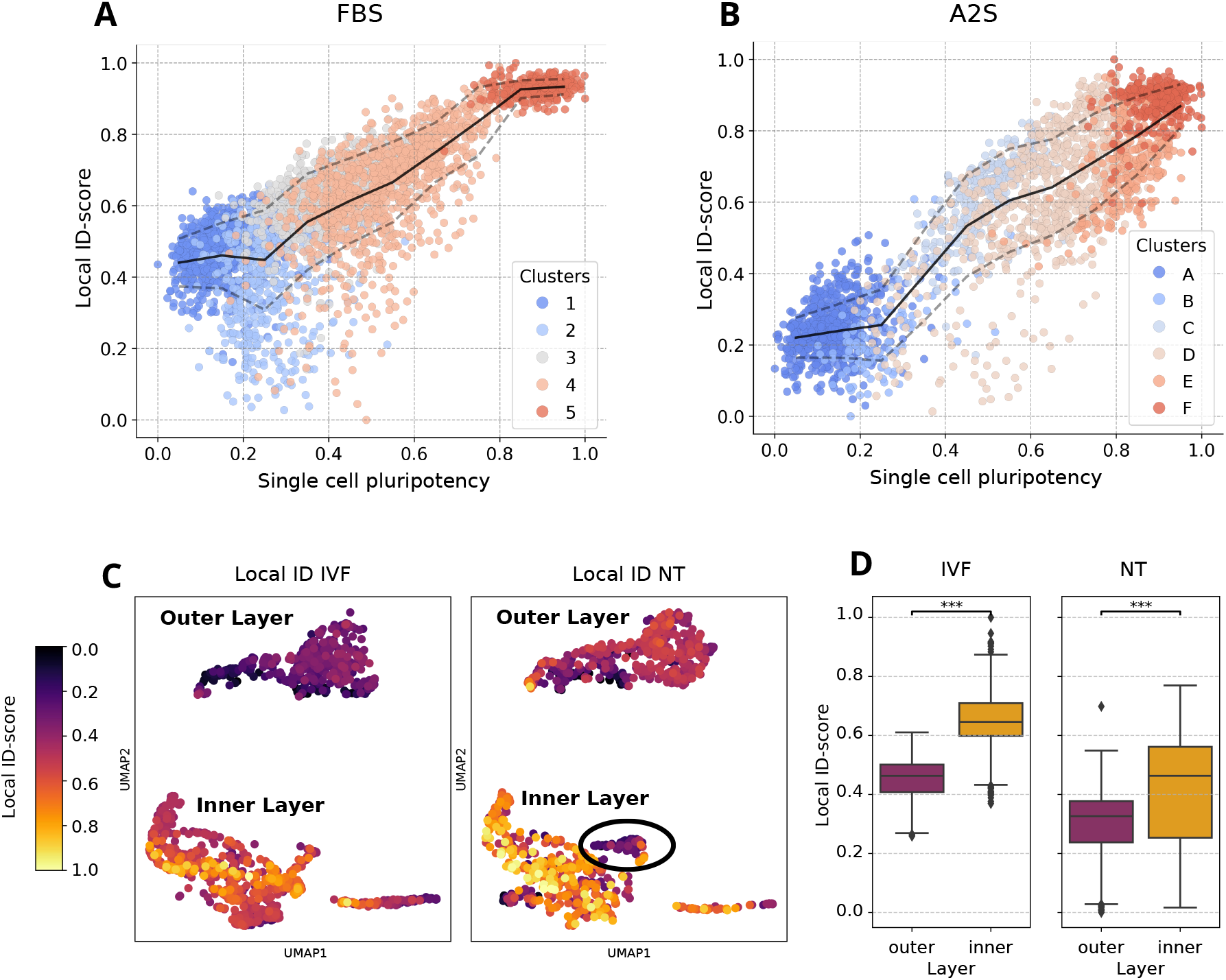
Local intrinsic dimension estimation as a single cell potency measure -. **A, B)** Scatter plots of local ID-scores vs pluripotency score for each cell in the FBS condition (panel A) and A2S condition (panel B) from dataset [49]. Each dot represents a single cell and is colored according to the cluster assignment. Pluripotency scores were calculated as described in Section 4.3.1. The continuous black line indicates the mean local ID-scores across binned values of pluripotency (number of bins = 20), dashed lines indicate the standard deviation. Local ID was estimated using 600 nearest neighbors. Pearson correlation coefficients are respectively 0.80 (panel A) and 0.89 (panel B). **C)** UMAP projections of data from [61] for the *in vitro* fertilization (IVF) data (left panel) and nuclear transfer (NT) data (right panel). Color encodes for the local ID-score for each cell computed using 100 neighbors. **D)** Boxplot of the local ID distributions of the inner and outer layer for the IVF condition (left panel) and NT condition (right panel).

We then extended our analysis to a recently published dataset derived from an *in vivo* nuclear reprogramming experiment in [61]. In this study, nuclei from endodermal cells were transferred into oocytes, which were subsequently fertilized to generate nuclear transfer (NT) embryos. Compared to *in vitro* fertilization (IVF)-derived control embryos, NT embryos frequently fail to develop. Previous analyses have shown that reprogramming efficiency depends on cell type, with ectodermal cells from the inner layer (e.g., basal stem cells) being particularly susceptible to reprogramming defects, which lead to impaired differentiation and disrupted body patterning [61]. We reanalyzed scRNA-seq data generated from the ectoderm of NT and IVF embryos [61]. These data were clustered and annotated using previously known marker genes, allowing the identification of two main groups of cells: one coming from the inner layer, which includes cells with generally higher potency (including basal stem cells) and another coming from the outer layer.

When we quantified differences in local ID between the two conditions (IVF vs NT embryos) and across the two layers, we observed that outer layer cells consistently exhibited lower ID values than inner layer cells in both IVF and NT embryos (Figure 5 C,D). This supports the notion that inner layer cells are generally less differentiated. Moreover, when comparing inner layer ID distributions between IVF and NT conditions, we detected a greater variability in the NT embryos. This finding aligns with prior reports that inner layer cells in NT embryos exhibit greater reprogramming-associated perturbations and differentiation potential defects [61]. Notably, the local ID analysis reveals the presence of an NT-specific group of cells in the inner layer, displaying an ID score lower than that of the surrounding cells (see the circled cells in Fig. 5C). This group of cells was reported to express an atypical sets of markers specific to ectodermal (e.g., *sox11*) and endodermal (e.g., *sox17b* and *foxa4*) fates, and was suggested to represent a terminal cell state produced by the disrupted differentiation dynamics in the inner-layer cells of the NT [61].

This analysis highlights several key advantages of the local ID estimation. First, it enables the distinction between inner and outer ectodermal layers in a completely unsupervised, marker-agnostic manner. Second, it reveals that the most significant perturbations in potency occur in inner layer cells, consistent with previous conclusions drawn from more supervised analyses and experimental validations [61].

Our findings demonstrate that the local ID estimation provides a powerful, unbiased framework for assessing cellular potency and heterogeneity at single-cell resolution. This continuous measure can also be exploited for gene discovery, by identifying genes whose expression significantly correlates with the ID. Supplementary Figure 5 illustrates an example of this analysis using the same dataset of Figure 5A,B. We computed the Pearson correlation between the expression of each gene and the local ID values in FBS and A2S conditions. The global distribution of expression–ID correlations is centered around zero, indicating no overall trend. However, when restricting the analysis to the set of differentiation-related genes used to define the pluripotency score, the distribution is significantly shifted toward negative correlation values. This shift is consistent with the positive association between the ID and the pluripotency score observed in Figure 5 A,B. Focusing specifically on transcription factors (TFs), we identified several TFs whose expression levels strongly correlate with the local ID. For example, *Utf1* and *Mtf2* consistently ranked as the top correlating TFs under both culture conditions. These genes are well-established regulators of pluripotency and have been implicated in the reprogramming of somatic cells into iPSCs [62, 63]. Additional TFs, such as *Pou5f1, Tcf15, Tfdp1*, and *E2f4*, recognized for their role in maintaining or modulating pluripotency [64, 65, 66, 67], have also emerged as highly correlated candidates.

## 3 Discussion

In this study, we show how quantifying the intrinsic dimensionality of scRNA-seq data offers a new lens through which cellular reprogramming can be explored. Unlike traditional analyses that rely on predefined marker genes and prior biological knowledge, our approach focuses on the global geometrical structure of the data, providing a marker-agnostic, unsupervised measure of cellular potency. We define an ID-score based on the intrinsic dimensionality and provide an open-source Python package, IDEAS, to facilitate its computation. We demonstrate that IDEAS is a powerful tool for quantifying cellular plasticity along reprogramming trajectories.

Reprogramming is a process with profound implications for regenerative medicine and our understanding of cell identity, yet remains difficult to study due to its inefficiency, heterogeneity, and the limitations of marker-based analysis. By applying IDEAS to scRNA-seq data from reprogramming experiments, we show that it effectively captures the progression of cells through pluripotent states. Specifically, ID-scores successfully align cell clusters along reprogramming trajectories in a manner consistent with supervised analyses based on established differentiation markers, yet without requiring any prior biological knowledge. The ID-score also proves useful in distinguishing between different lineage trajectories, even in complex branching scenarios. For example, in datasets where reprogramming and differentiation occur simultaneously and lead to multiple cellular identities, the ID-score provides a clear, unsupervised separation of these divergent paths.

Importantly, we extend this analysis to the single-cell level by localizing the computation of ID to individual cell neighborhoods, enabling exploration of potency variability within and across clusters. This fine-grained resolution allows for the detection of subpopulations and subtle identity shifts that would be missed by coarse-grained or marker-dependent methods. Notably, we apply this approach to an *in vivo* reprogramming dataset and demonstrate that the ID-score uncovers complex features of the epigenetic landscape. The single-cell ID-score also opens up new opportunities for gene discovery by identifying genes whose expression is significantly correlated with the ID. Using this approach in an illustrative example, we identified key transcription factors such as *Utf1* and *Mtf2*, often associated with high-potency states, together other known pluripotency regulators. Therefore, the ID score could represent a tool for uncovering new molecular drivers of pluripotency and reprogramming.

This work underscores the broader utility of analyzing global properties of scRNA-seq data to fully exploit its high-dimensional nature. Drawing an analogy with statistical physics, we can think of the ID-based potency score as a thermodynamic observable that reflects the collective cellular state, rather than focusing on potentially noisy, gene-specific signals. While the ID-score does not rely on prior biological knowledge and marker-based analyses, it can be used to complement them: combining unsupervised, geometry-based analyses like the ID-score with known markers or annotations can enrich biological interpretation and support more comprehensive modeling of cell fate changes.

Although our primary focus was on cellular reprogramming, the ID analysis also shows promise for other biological processes involving dynamic changes in cell potency, such as embryonic development [22], cancer, immune activation, and cellular aging. Another possible extension could be the integration of additional single-cell omics modalities (e.g., ATAC-seq, CITE-seq), which may reveal distinct global geometrical signatures across layers of regulation.

To further improve the robustness of the ID-score, several limitations should be addressed in future work. The estimation of intrinsic dimensionality is sensitive to the choice of distance metric in gene expression space. In the ~10,000-dimensional gene space, conventional Euclidean distance is not necessarily the optimal choice. Alternative similarity measures, such as cosine similarity or Manhattan distance, could be explored and incorporated as options within the IDEAS framework.

In conclusion, the ID-score can become a valuable addition to the single-cell analysis toolkit. Thanks to its flexibility, interpretability, and marker-independent formulation, it offers a simple computational method to infer cell potency and can complement well-established approaches such as trajectory inference and pseudotime analysis, while also enabling the discovery of potency-associated genes. Additional benchmarking and applications to a broader range of biological systems will be critical next steps in establishing the full potential of this approach.

## 4 Methods

### 4.1 Datasets information

The scRNA-seq datasets collected for this study are freely accessible through the GEO repositories indicated below [68].

#### 4.1.1 Dataset 1 : [49]

##### Raw data available at GEO: GSE108222

In [49], mouse embryonic fibroblasts (MEFs) were reprogrammed into induced pluripotent stem cells (iPSCs) via the expression of OSKM factors (Oct4, Sox2, Klf4, c-Myc) under two culture conditions: FBS (Fetal Bovine Serum) and A2S (ascorbic acid, 2i, SGC), the latter designed to enhance reprogramming efficiency. The raw count matrix included 8,334 cells and 24,421 genes, sequenced at multiple time points (FBS: days 0, 3, 6, 9, 12; A2S: days 0, 2, 4, 6; and final mESCs). Building on the observations and analytical pipeline described in [49], ESCs cultured in FBS were selected as the endpoint for both experiments, as their gene expression profiles most closely resembled those of reprogrammed cells under both conditions [49].The resulting datasets included 3,435 cells (FBS) and 2,408 cells (A2S). Before intrinsic dimension (ID) analysis with IDEAS, we annotated each cell with metadata including clustering, pluripotency score, and pseudotime inference (detailed in Sections 4.3). For the ID estimation, we used the raw count matrix with the following filtering: we excluded genes expressed in fewer than 1% of cells and retained only protein-coding genes, resulting in 11,776 genes for the FBS dataset and 12,193 for A2S.

#### 4.1.2 Dataset 2 : [59]

##### Data available at GEO: GSE112004

In [59] an initial population of pre-B cells was induced to either transdifferentiate into macrophages or reprogram into iPSCs. In both protocols, cells were first incubated with beta-estradiol for 18h to activate C/EBP*α* (collecting cells at 0 h, 6 h, 18 h), generating a “poised state”. Macrophage differentiation continued with extended C/EBP*α* exposure (samples at 42 h, 66 h, 114 h), while reprogramming was initiated via OSKM induction (samples at days 2, 4, 6, 8). Two replicates of 192 cells were collected per time point. The raw scRNA-seq count matrix was constructed by concatenating all time points and conditions, retaining only genes common to all samples. This yielded 11,838 genes ×1,536 cells for reprogramming and 11,838 genes ×1,152 cells for transdifferentiation. All genes passed filtering (expressed in *≥* 1% of cells).

#### 4.1.3 Dataset 3 : [34]

##### Data available at GEO:GSE242424

In [34] reprogramming from human embryonic fibroblast to iPSCs is performed through OSKM induction. The scRNA-seq data were collected at days 0, 2, 4, 6, 8, 10, 12, 14 and the endpoint of iPSCs at passage 30. Processed data produced by the authors of [34], including count matrix, quality-controlled barcodes, gene list, cluster annotations, and UMAP coordinates, are available at Zenodo. The scRNA-seq data yielded 59,378 high-quality cells ×29,165 detected genes. After filtering out genes expressed in fewer than 1% of cells, 14,889 genes were retained. The metadata includes 15 clusters defined using the Leiden algorithm. Cluster 2 (140 cells) was excluded from ID analysis due to low cell count, resulting in 14 clusters ranging from 523 cells (cluster 7) to 11,041 cells (cluster 1).

#### 4.1.4 Dataset 4 : [61]

##### Data available at GEO: GSE269252

In [61], nuclei from endodermal cells were transferred into enucleated *Xenopus laevis* eggs to generate nuclear transfer (NT) embryos. The differentiation of the produced reprogrammed cells into epidermis was compared to *in vitro* fertilization (IVF)-derived controls. Single-cell RNA-seq was performed at gastrula stage 12 across two experiments, yielding 1,841 IVF and 1,564 NT cells. For ID analysis, we selected one batch per condition with the highest cell counts (SIGAH5 (IVF) and SIGAH12 (NT)) resulting in 1,267 IVF and 1,043 NT cells.

### 4.2 ID-score computation

Given *G* cellular populations (different cell-types/developmental stages/patients…) and *S* equally represented sub-samplings of each of them, IDEAS performs the ID estimation *G · S* times. These ID values returned as output can be rescaled to the [0,1] range by setting the `id_score` parameter to *True*. The rescaling procedure is defined on single ID measurements as:

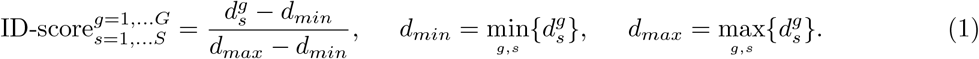

If IDEAS is used with `full_output` set to *False*, and `id_score` set to *True*, the mean and standard deviation across groups are computed after the rescaling.

### 4.3 Metadata generation

Metadata annotations for the dataset from [49] were generated using the Scanpy library [23]. The resulting annotations include, for each single cell: sampling day, biological replicate, culture condition (FBS or A2S), cluster identity, pluripotency score, pseudotime value, and UMAP coordinates. All metadata are contained in the metadata files available at maddalenastn/IDEAS on GitHub.

#### 4.3.1 Pluripotency score

A pluripotency score was calculated for each of the *N* cells using the score genes function implemented in the Scanpy package [23]. This score *s*_*i*_ (with *i* = 1, 2, … *N*) corresponds to the difference between the average expression of a particular set of genes and a reference set of genes, randomly sampled for each binned expression value.

In the case of mouse embryonic fibroblasts reprogramming to iPSC, the set of genes is given by the list of differentiation-related genes identified in [49] (Table S1, groups A,B,C,D) by differential expression analysis and Gene Ontology annotation. Assuming that pluripotent cells underexpress genes associated with differentiation, a pluripotency score at single-cell level is formulated as:

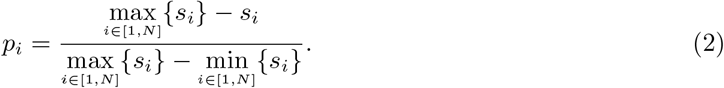

The obtained score ranges between 0 (least pluripotent, most differentiated) and 1 (most pluripotent, least differentiated). To associate a pluripotency score with a whole cluster (Fig. 2), the average pluripotency score across the cells of that cluster is computed, taking the standard deviation as error.

#### 4.3.2 Clustering

Clustering was performed using the Scanpy library [23]. Cell counts were normalized using total-count normalization (`target_sum=1e6`), log-transformed (`sc.pp.log1p`), and highly variable genes (HVG) were selected (`min_mean=0.0125`, `max_mean=3`, `min_disp=0.5`). The dataset was then split by culture condition (FBS and A2S) and analyzed separately. The first 50 principal components were computed applying the PCA method on the highly variable genes using the ‘`arpack`‘solver. The neighborhood graph was constructed with the `correlation` distance metric, and clustering was performed via the Leiden algorithm [55] with a resolution of 0.5. UMAP [56] was computed for 2D visualization. Clusters containing less than 1% of total cells were excluded from downstream analysis resulting in 5 clusters for FBS and 6 for A2S. The two biological replicates were uniformly represented across all clusters. The pluripotency score was computed for each cell as described in section 4.3.1, and clusters were ordered and renamed according to their mean pluripotency levels.

#### 4.3.3 Pseudotime Analysis

Pseudotime analysis was conducted separately for the FBS and A2S datasets, after performing the same normalization, log-transformation and HVG selection described in 4.3.2. The neighborhood graph was then created using `n_neighbors = 20`, using the ‘`X`‘representation and the ‘`gauss`‘method. The first 15 diffusion components were obtained [69] and used to compute the diffusion pseudotime, using one of the MEF cells as the root (starting point) of the trajectory. The analysis produced a single pseudotime value for each cell, normalized between 0 and 1. To compute a single score for each cluster, we averaged the pseudotime across all cells within the cluster, taking the standard deviation as the error.

### 4.4 Intrinsic dimension estimators

IDEAS allows the user to specify the estimator for the intrinsic dimension. The choice is between TWO-NN and a PCA-based observable. The first is a fractal estimator that leverages the statistics of the distance of each point from its first two neighbors [43]. The second corresponds to the number of principal components that are necessary to retain a given percentage of the data variance (variance_e parameter). A formal definition of these two quantities is in the Section 7.1 of Supplementary Material.

Although PCA is intuitive, it relies on a linear transformation, making it suitable mostly for data that is linearly embedded. In contrast, TWO-NN can capture non-linear structures due to its local nature. Furthermore, it is less sensitive to undersampling than projective methods like PCA, which typically require an exponentially large number of points relative to the original embedding dimension to yield accurate estimates. Finally, as shown by Biondo et al. [22], the computation of principal components can be strongly affected by outliers or by the presence of multiple clusters within the data.

For these reasons, we present only the results obtained with TWO-NN in the main text. However, compatible results can be obtained using a PCA-based estimator (see Methods).

## 5 Code and data availability

All code needed to run IDEAS is available at maddalenastn/IDEAS on GitHub. This repository contains instructions for the installation and usage of the IDEAS tool and Jupyter notebooks to reproduce the results in all figures. All scRNA-seq datasets are publicly accessible in the GEO repository through the corresponding GEO accession numbers reported in the Methods section 4.1.

## 6 Acknowledgments

We would like to thank Giacomo Masserdotti, and the members of the Scialdone and Osella labs for the constructive feedback, insightful discussions, and helpful comments. This work was supported by the Helmholtz Association (A.S.), the Deutsche Forschungsgemeinschaft (Project number 448727785 to A.S.), and by the Italian “Ministero dell’Università e della Ricerca”, PRIN 2022 - COD. 2022PY8MHN - GeCoS: Genomic Component Systems.

Niccolò Cirone is a PhD student enrolled in the National PhD program in Artificial Intelligence, XXXIX cycle, course on Health and life sciences, organized by Università Campus Bio-Medico di Roma.

Analyzed the data: M.S., N.C., M.B. Wrote the manuscript: M.S., N.C., M.B., M.O., A.S. Conceived and supervised the project: M.O., A.S. All authors approved the final draft of the manuscript.

