## Supplementary Material for "Intrinsic dimensionality of single-cell transcriptomic data reveals potency landscapes during cell reprogramming"

#### 1 Intrinsic dimension estimators

##### 1.1 Two Nearest Neighbors

The Two Nearest Neighbors (TWO-NN) algorithm has been proposed in [1] as an intrinsic dimension estimator based exclusively on local properties of the data. In particular, the algorithm infers the ID from the statistics of the first and second nearest neighbors of each point.

This method assumes that the $N$ data points $\mathbf{x_1}, \mathbf{x_2}, \ldots, \mathbf{x_N}$ are independent and identically distributed samples from a probability density $\rho(x)$ whose support is a $d$-dimensional manifold, where $d$ is the intrinsic dimension we aim to estimate. The second main assumption of the model is that $\rho(x)$ is locally uniform, i.e., approximately constant in the spherical neighborhood centered at $x_i$ and radius given by the distance with its second neighbor.

Reference [1] shows that for a given data point $\mathbf{x_i}$, the ratio $\mu_i := \frac{r_i^{(2)}}{r_i^{(1)}}$ of its second nearest neighbor distance ($r_i^{(2)}$) to its first nearest neighbor distance ($r_i^{(1)}$) follows a Pareto distribution:

$$f(\mu_i|d) = d\mu_i^{-d-1}. \tag{1}$$

The cumulative distribution is then obtained by integration as:

$$F(\mu_i|d) = 1 - \mu_i^{-d}. \tag{2}$$

Sorting the values of $\mu_i$ in ascending order, the empirical cumulative distribution is given by $F^{emp}(\mu_i) \doteq \frac{i}{N}$. This means that, knowing the set of $\mu_i$ for different data points, $d$ can be computed as the slope of the line that fits the points of coordinates $\{(\log(\mu_i),\ (-\log(1 - F^{emp}(\mu_i))) \mid i = 1, ..., N\}$.

This procedure is significantly affected by a few points with high values of $\mu$, which make the estimation potentially unstable. To solve this issue and gain robustness, the authors propose excluding from the regression points that fall above the 90th percentile.

##### 1.2 Principal Component Analysis

Principal Component Analysis (PCA) is a well known dimensionality reduction technique based on a linear projection on a lower dimensional space. The new set of coordinates represent the "most relevant features" of the system, along which data points display higher variability.

Given N data points (cells), and m features (genes), the covariance matrix C is a symmetrical $mxm$ matrix. From a mathematical point of view, PCA relies on the diagonalization of C, with eigenvectors (principal components) being the axes of the new coordinate system. These can be ranked according to the magnitude of their corresponding eigenvalues $\lambda_1, \lambda_2, ..., \lambda_m$, that coincide with the variance retained by each component.

The sum of all eigenvalues corresponds to the total data variance V:

$$\sum_{i=1}^{m} \lambda_i = V, \tag{3}$$

and the shape of their rank-plot encodes information about the intrinsic dimension $d$ of the dataset. Specifically, a common procedure consists in counting how many components are needed to retain a specific percentage $f$ of the data variance:

$$d = \max_{\widetilde{d} \in \mathbb{Z}^+} \left\{ \widetilde{d} : \sum_{i=1}^{\widetilde{d}} \lambda_i \geq fV \right\}. \tag{4}$$

Although very intuitive, PCA-based estimators leverage a linear transformation, making them reliable only in case of linearly embedded data. In case of curved manifolds, Eq. (4) leads to an overestimation of the intrinsic dimension.

### 2 Technical properties of ID : sensitivity to data heterogeneity

Disentangling the effects due to different sources of cellular variability represents one of the most typical and challenging issues in scRNA-seq analysis. In time-course experiments, grouping cells from different days may result in a mixture of distinct cell types present on the same time point, as discussed in Main Section 2.1. Although unsupervised methods like clustering are often used to address cell-type mixing, they do not ensure unambiguous cell identities due to their dependence on resolution and potential imbalances in cell-type abundance. In this section, we focus on how the coarse-grained structure of single-cell data affects the intrinsic dimension estimation, and propose the local sampling as a possible strategy to tackle this issue.

The case study is represented by a filtered version of the dataset [2], including only 3000 randomly sampled cells belonging to fibroblasts ($\mathcal{F}$) and iPSCs ($\mathcal{P}$), respectively cluster 1 and 15 in Fig.4 (Supplementary Fig. 1A). In Supplementary Fig. 1B, the ID-score obtained separately on $\mathcal{F}$ and $\mathcal{P}$ is reported, as well as the value corresponding to the union $\mathcal{F}$ U $\mathcal{P}$. Interestingly, the ID for the combined set is closer to the ID of $\mathcal{F}$ than to the one of $\mathcal{P}$. This result suggests that data points from low-dimensional manifolds may have a higher impact on the overall ID estimation with respect to those from high-dimensional manifolds, as suggested by Biondo et al. [3].

A quantitative study of this effect as a function of cell type heterogeneity is shown in Supplementary Fig. 1C. The x-axis gives the percentage of cells from cluster $\mathcal{F}$ on the total of 3000 cells sampled from the composite manifold $\mathcal{F}$ U $\mathcal{P}$. At each iteration, an exponentially increasing number of randomly sampled cells from $\mathcal{F}$ is added, while removing the same number of cells from $\mathcal{P}$, in order to maintain the same number of points across iterations. When the proportion is 99 to 1, the effects due to the co-presence of two cell types are already clear. The magnitude of this ID drop mainly depends on the difference of intrinsic dimension between the mixed populations [3].

These results suggest that a single value of ID is insufficient to fully characterize a heterogeneous cellular population. This poses a challenge, especially since in most cases we aim to estimate the population's ID without prior knowledge of its internal variability. A more effective strategy is to adopt a local sampling approach, such as the *local_2nn* method. By assigning an ID value to each individual cell based on its local neighborhood, the composition of the whole population in terms of ID can be analyzed, giving insights on the granular nature of the data. Applying this method to the combined population $\mathcal{F}$ U $\mathcal{P}$ yielded a bimodal ID distribution (Fig. 1D), reflecting the dataset's bipartite structure. Additionally, the spread of ID values within each group provided insight into internal variability, with fibroblasts exhibiting higher dispersion in cell potency compared to iPSCs.

Taken all together, these results show a case study where adopting the *local_2nn* method can give insights on the data that would be otherwise hidden in global ID estimates, revealing subpopulation structures and cellular heterogeneity.

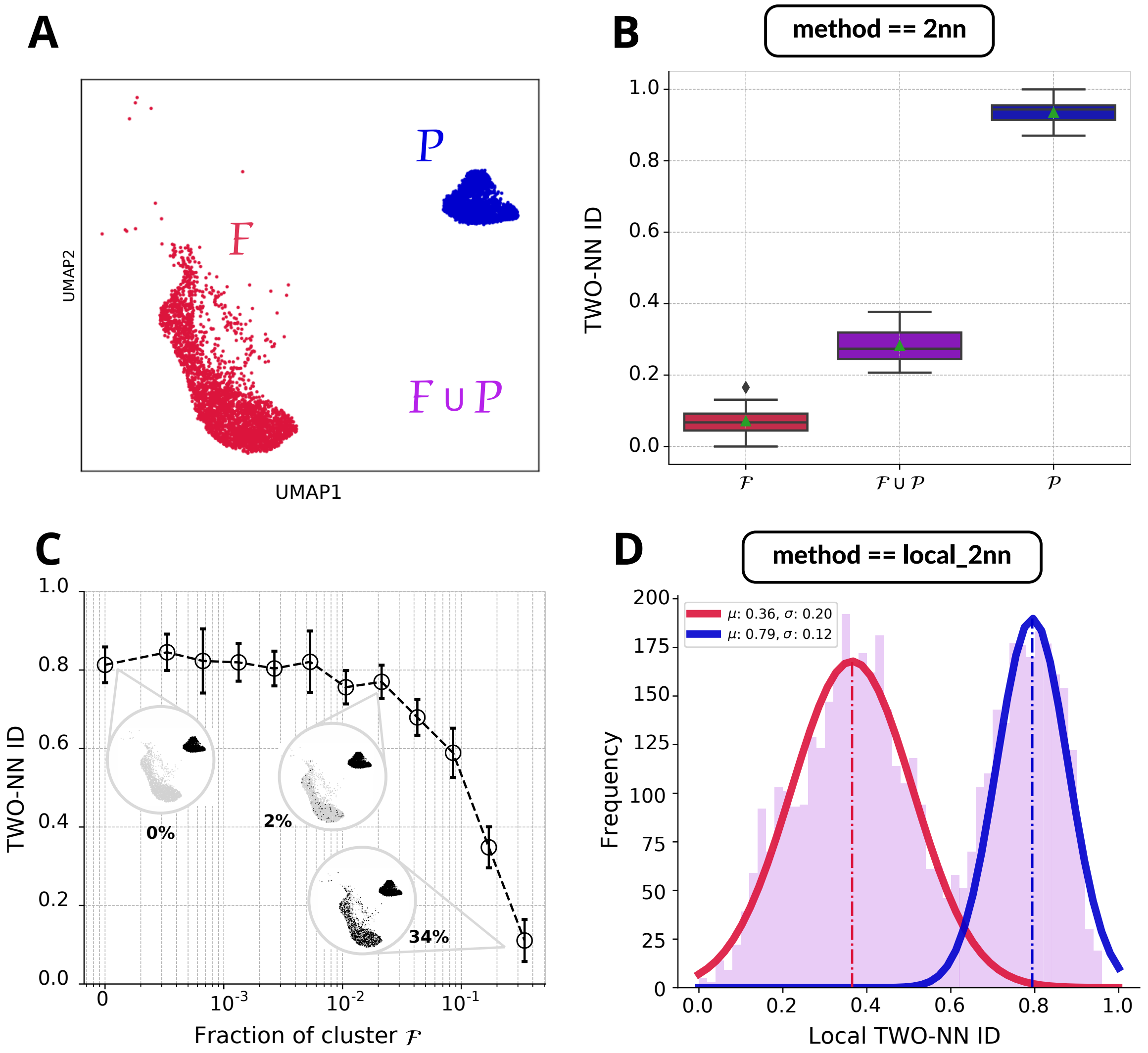


Supplementary Figure 1: **Sensitivity of ID to data heterogeneity: A)** UMAP projection of fibroblasts cluster ($\mathcal{F}$) in red and iPSCs cluster ($\mathcal{P}$) in blue. Each cluster is composed of 3000 randomly sampled cells from cluster 1 and cluster 15 in dataset [2]. In purple we denote the union of the two ($\mathcal{F} \cup \mathcal{P}$), which includes a total number of cells equal to 6000. **- B)** Boxplot of the ID estimations obtained respectively on $\mathcal{F}$, on $\mathcal{F} \cup \mathcal{P}$ and on $\mathcal{P}$ through the TWO-NN method. The sample size was set to 2400 and the measure was repeated over 30 samples. **- C)** Plot of the ID obtained via the TWO-NN method as a function of cell type heterogeneity. Cell type heterogeneity is varied by sampling 3000 cells from the composite manifold F U P, starting with only cells from cluster P and gradually increasing the proportion of cells from cluster F, while keeping the total sample size constant to 3000. x-axis indicates the fraction of cells from cluster F in each sampling. For each fraction, the ID is estimated 10 times; the plot shows the mean ID (dots) and standard deviation (error bars) across ID measurements. **- D)** Histogram of the ID measurements obtained with the local TWO-NN method on $\mathcal{F} \cup \mathcal{P}$ . The number of neighbors was set to 600, and the number of samples to 5000. The histogram reveals a bimodal distribution that can be fitted with two gaussian curves and reveal the bipartite structure of the data.

### 3 Pluripotency score and pseudotime ordering

Inferring an ordering of clusters or single cells based on their potency level is a complex problem that can be approached in multiple ways. In Section 2.1.1, we proposed an ordering based on a pluripotency score defined using the expression levels of genes associated with cellular differentiation, as determined by their Gene Ontology annotations. This represents a supervised approach that incorporates prior biological knowledge about gene function. As an orthogonal strategy, we also considered pseudotime inference, which orders cells along a trajectory based on transcriptional similarity, given a predefined starting point (see Methods section). To validate the ordering obtained via the pluripotency score, we compared it with the pseudotime-derived ordering. Supplementary Figs. 2A,C show the correlation between the pluripotency score and pseudotime across clusters in dataset [4], indicating that both methods yield a consistent cluster ordering both in the FBS and A2S experiments. A similarly strong positive correlation is observed between pluripotency score and pseudotime at the single-cell level (Supplementary Figs. 2B,D). Together, these results indicate a clear concordance between pseudotime and pluripotency score ordering, both at the cluster and single-cell resolution. Finally, we evaluated whether local ID increases with pseudotime, as observed with the pluripotency score in Fig. 5A,B, and confirmed the presence of a positive correlation (Supplementary Fig. 3).

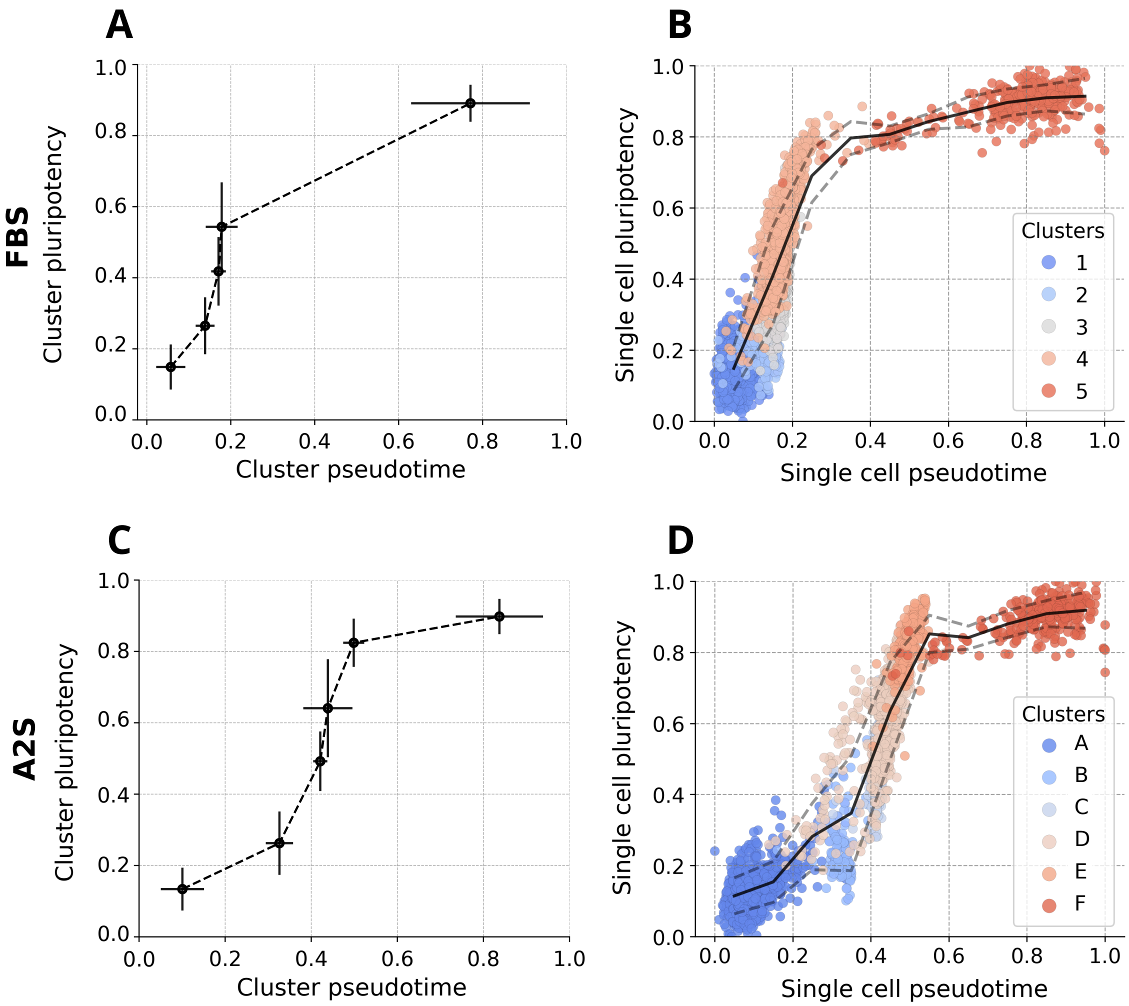


Supplementary Figure 2: **Pseudotime and pluripotency score - A,C)** Mean pluripotency score over clusters versus mean pseudotime over clusters for the FBS (panel A) and A2S (panel C) conditions [4]. Pluripotency and pseudotime on single cells were estimated as explained in the Methods section. Error bars represent standard deviation of pluripotency (vertical error bars) and pseudotime (horizontal error bars) across cells from each cluster. **- B, D)** Scatter plot of single cell pluripotency versus single cell pseudotime for the FBS (panel B) and A2S (panel D) conditions. Each dot represents a single cell and is colored according to its cluster. The continuous black line represents the trend of the mean single cell pluripotency across binned values of pseudotime (number of bins = 10), the dashed lines indicate the standard deviation of pluripotency values.

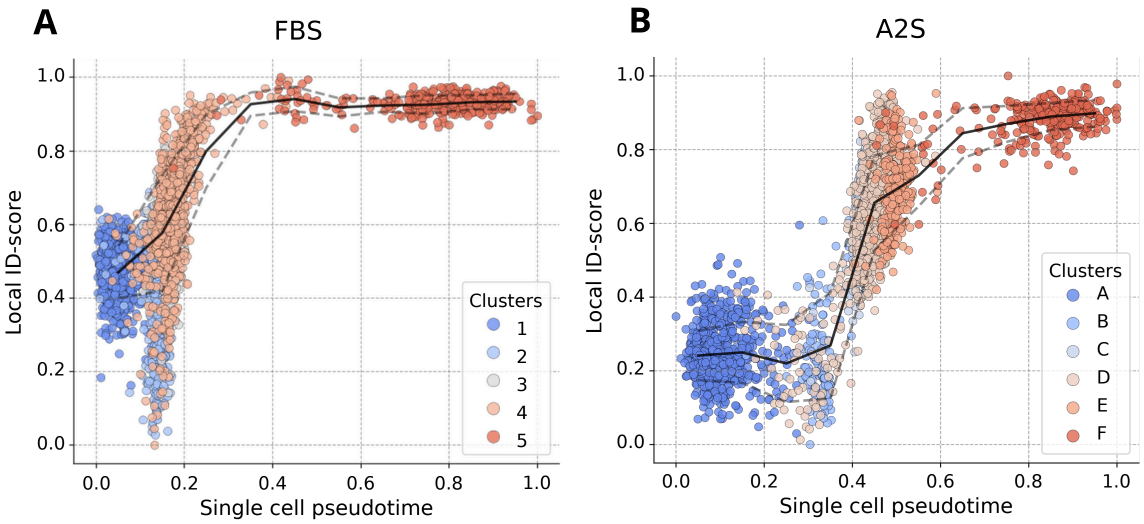


Supplementary Figure 3: **Local Intrinsic dimension correlates with pseudotime -** Scatter plots of local intrinsic dimension (Local ID) values vs pseudotime for each cell in the FBS condition (panel **A)**) and A2S condition (panel **B)**) from dataset [4]. Each dot represents a single cell and is colored according to the cluster assignment. Pseudotime is calculated as described in the Methods section. The continuous black line indicates the mean local ID across binned values of pseudotime (number of bins = 10), dashed lines indicate the standard deviation of local ID values. Local intrinsic dimension was estimated using 600 nearest neighbors.

#### 4 Local Intrinsic Dimension resolution

In Fig. 5A,B, we show the positive correlation existing between the single-cell version of the intrinsic dimension and the gene-informed score defined in the Methods section, interpretable as the ground truth for cell pluripotency. The local ID of a cell corresponds to the ID-score relative to the neighborhood of that cell in the gene expression space. In the following, we outline two key considerations regarding the choice of the neighborhood size. First, from a technical perspective, small neighborhood sizes are more susceptible to noise [5], whereas too large sizes tend to capture global patterns, such as the coexistence of multiple cell types [3]). Second, biological systems are inherently hierarchical, often exhibiting a clustered organization [6]. The resulting distribution of cluster sizes introduces a typical scale that is generally unknown and can vary across systems. To ensure that our results are not sensitive to a particular neighborhood size choice, we studied how varying the number of neighbors affects the Pearson correlation between the local ID estimates and the pluripotency score. As shown in Supplementary Fig. 4, a broad range of neighborhood sizes yields high correlations ($> 0.6$).

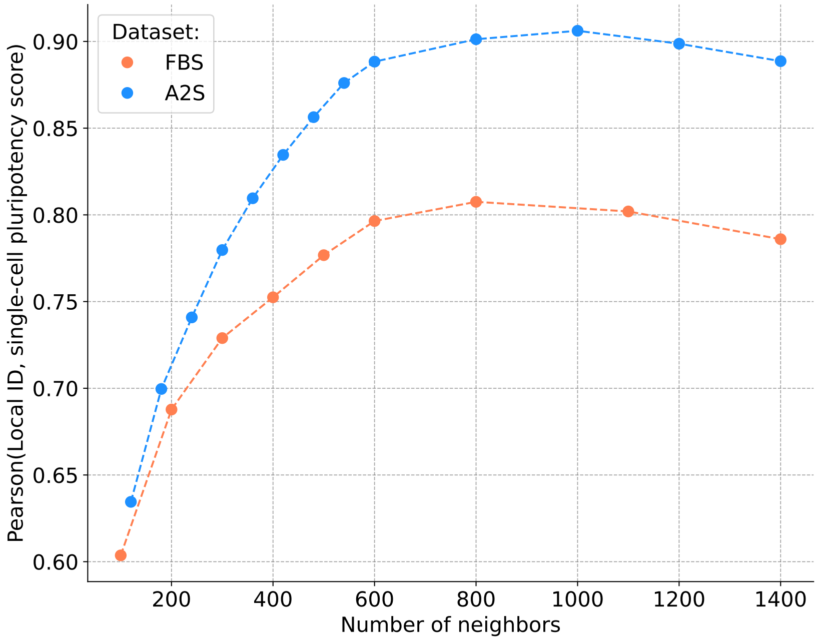


Supplementary Figure 4: **Local ID dependence on the number of neighbors.** Pearson correlation between the single-cell pluripotency score (see Methods) and the local ID calculated on different sizes of the neighborhood. The trend is shown both in FBS and A2S condition of iPSC generation dataset [4].

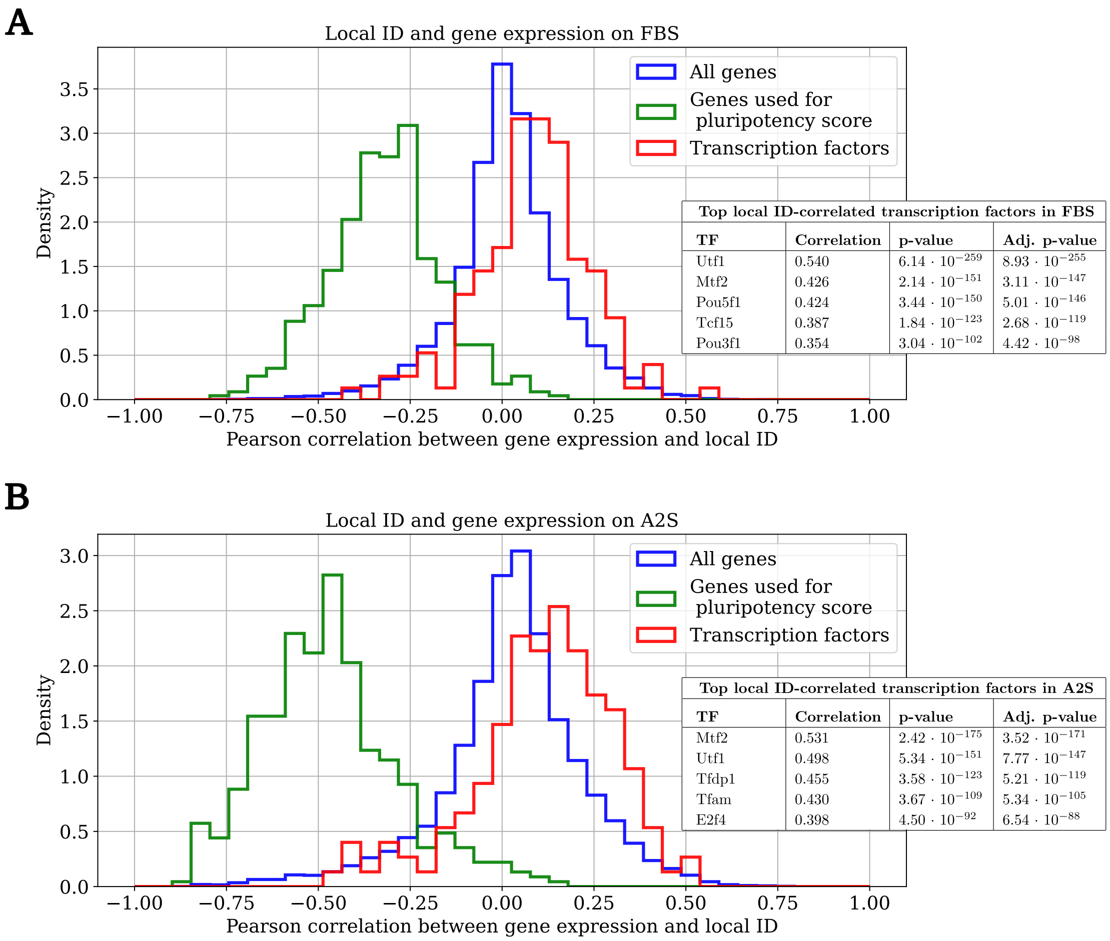


**Top local ID-correlated transcription factors in FBS**

| **TF** | **Correlation** | **p-value** | **Adj. p-value** |
|---|---|---|---|
| Utf1 | 0.540 | $6.14 \cdot 10^{-259}$ | $8.93 \cdot 10^{-255}$ |
| Mtf2 | 0.426 | $2.14 \cdot 10^{-151}$ | $3.11 \cdot 10^{-147}$ |
| Pou5f1 | 0.424 | $3.44 \cdot 10^{-150}$ | $5.01 \cdot 10^{-146}$ |
| Tcf15 | 0.387 | $1.84 \cdot 10^{-123}$ | $2.68 \cdot 10^{-119}$ |
| Pou3f1 | 0.354 | $3.04 \cdot 10^{-102}$ | $4.42 \cdot 10^{-98}$ |

**Top local ID-correlated transcription factors in A2S**

| **TF** | **Correlation** | **p-value** | **Adj. p-value** |
|---|---|---|---|
| Mtf2 | 0.531 | $2.42 \cdot 10^{-175}$ | $3.52 \cdot 10^{-171}$ |
| Utf1 | 0.498 | $5.34 \cdot 10^{-151}$ | $7.77 \cdot 10^{-147}$ |
| Tfdp1 | 0.455 | $3.58 \cdot 10^{-123}$ | $5.21 \cdot 10^{-119}$ |
| Tfam | 0.430 | $3.67 \cdot 10^{-109}$ | $5.34 \cdot 10^{-105}$ |
| E2f4 | 0.398 | $4.50 \cdot 10^{-92}$ | $6.54 \cdot 10^{-88}$ |

Supplementary Figure 5: **Correlation between local ID and single-gene expression.** Pearson correlation between the raw expression of single genes and the local ID across cells reprogrammed into iPCSs [4] is computed, considering two different culture conditions (fetal bovine serum - FBS - in **A**, ascorbic acid, 2i, SGC - A2S - in **B**). Blue curves refer to all the 14546 protein-coding genes sampled during the sequencing experiment. Green curves take in consideration only the differentiation-related genes used to formulate the pluripotency score (see Methods). Red curves represent the correlation distribution over mouse transcription factors (TFs) [7]. On the right, inset tables resume the top 5 TFs whose expression shows higher Pearson correlation with the local ID. Adjusted p-values correspond to p-values with Bonferroni correction, to account for multiple comparisons.
